# Desiccation-induced fibrous condensation of CAHS protein from an anhydrobiotic tardigrade

**DOI:** 10.1101/2021.06.22.449423

**Authors:** Maho Yagi-Utsumi, Kazuhiro Aoki, Hiroki Watanabe, Chihong Song, Seiji Nishimura, Tadashi Satoh, Saeko Yanaka, Christian Ganser, Sae Tanaka, Vincent Schnapka, Ean Wai Goh, Yuji Furutani, Kazuyoshi Murata, Takayuki Uchihashi, Kazuharu Arakawa, Koichi Kato

## Abstract

Anhydrobiosis is one of the most extensively studied forms of cryptobiosis that is induced in certain organisms as a response to desiccation. Anhydrobiotic species has been hypothesized to produce substances that can protect their biological components and/or cell membranes without water. In extremotolerant tardigrades, highly hydrophilic and heat-soluble protein families, cytosolic abundant heat-soluble (CAHS) proteins, have been identified, which are postulated to be integral parts of the tardigrades’ response to desiccation. However, the molecular mechanisms underlying these protein functions remain to be fully elucidated. In this study, we perfomed *in vitro* and *in vivo* characterizations of the self-assembling property of CAHS1 protein, a major isoform of CAHS proteins from *Ramazzottius varieornatus*, using a series of spectroscopic and microscopic techniques. Our *in vitro* observations showed that CAHS1 proteins homo-oligomerized *via* the *C*-terminal α-helical region and formed a hydrogel as their concentration increased, and that these molecular assembling processes were reversible. Furthermore, our *in vivo* observations demonstrated that the overexpressed CAHS1 proteins formed condensates under desiccation-mimicking conditions. These data strongly suggested that, upon drying, the CAHS1 proteins form oligomers and eventually underwent sol-gel transition in tardigrade cytosols. Thus, it is proposed that the CAHS1 proteins form the cytosolic fibrous condensates, which presumably have variable mechanisms for the desiccation tolerance of tardigrades. These findings provide insights into the protective mechanisms involved in the anhydrobiosis of tardigrades.

## Introduction

Anhydrobiosis is one of the most extensively studied forms of cryptobiosis that is induced in response to desiccation conditions in certain organisms, including bacteria, fungi, plants, and animals (1–4). Even after almost complete dehydration, anhydrobiotic organisms do not exhibit irreversible protein denaturation and aggregation. The organisms in anhydrobiotic states can tolerate desiccation for decades while maintaining their resuscitation ability (5). In this state, they are also cross-tolerant to various environmental stresses caused by extremely high and low temperatures, high and low pressures, and radiation (6–8). Therefore, a deeper understanding of the mechanisms behind the anhydrobiosis has versatile applications to develop technologies to endow molecular systems, cells, and tissues of clinical and industrial interest with resistance against various environmental stresses.

It has been hypothesized that anhydrobiotic species accumulate substances which can protect their biological components or cell membrane without water. Early studies on desiccation tolerance identified the disaccharide trehalose and the late embryogenesis abundant (LEA) proteins as the functional mediators of anhydrobiosis (9). Trehalose is known to accumulate at high levels under desiccating conditions in many anhydrobiotic organisms, including the larvae of sleeping chironomids *Polypedilum vanderplanki*, cysts of the brine shrimp *Artemia*, and vegetative tissues of the desert resurrection plant *Selaginella* (10–14). Trehalose is assumed to exert its protective functions through water replacement and vitrification; these are two distinct but not mutually exclusive mechanisms in protecting the organisms from the harmful effects of desiccation (12, 15). In the former mechanism, accumulating trehalose molecules extensively interact with biomolecular surfaces through hydrogen bonds by replacing the water molecules. In the latter mechanism, an amorphous state of trehalose without water vitrifies at a high concentration, preventing the movement of biomolecules, and thereby physically suppresses their denaturation. LEA proteins are a group of intrinsically disordered proteins originally discovered in plant seeds. They have been recently linked to anhydrobiosis in animals, such as nematodes, bdelloid rotifers, and crustaceans (3, 9, 16, 17). LEA proteins commonly share highly hydrophilic and heat-soluble properties and are supposed to alter their structures from a disordered state to an α-helical state under anhydrous conditions. Hence, LEA proteins have been related to desiccation tolerance with several hypothetical functions, including the stabilization of trehalose vitrification and molecular shielding to prevent protein denaturation and aggregation, regulation of the drying rate as a hydration buffer, and scavenging of divalent metal ions or reactive oxygen species (3, 9, 18).

In extremotolerant tardigrades, through heat-soluble proteomics in a search for LEA and LEA-like proteins, highly hydrophilic and heat-soluble protein families have been identified as cytosolic abundant heat-soluble (CAHS) proteins (19–22). Although these proteins are distinct from LEA proteins based on their conserved sequences, they share some similarities with LEA proteins in terms of the bioinformatic prediction of high propensity for α-helix formation (19, 20). Since neither the LEA proteins nor trehalose are abundant in tardigrades, these tardigrade-specific abundant proteins are postulated to be integral parts of their response to desiccation as alternatives. While CAHS proteins are constitutively and abundantly expressed in *Ramazzottius varieornatus*, they are strongly induced under desiccation stress in another tardigrade species *Hypsibius exemplaris;* this induction requires *de novo* gene expression and is indispensable for survival (23, 24). This was subsequently confirmed by RNAi experiments and by the improved desiccation tolerance of yeast and bacterial cells with the heterologous expression of CAHS proteins (21). Thus, this growing evidence suggests that the CAHS and LEA proteins play crucial roles in anhydrobiotic responses. However, the molecular mechanisms underlying these protein functions remain to be fully elucidated.

In view of the situation, we herein conducted *in vitro* and *in vivo* characterizations of molecular assembly of CAHS1, a major isoform of CAHS proteins from *R. varieornatus*, using a series of spectroscopic and microscopic techniques. Our observations revealed that the CAHS1 proteins self-assembled into α-helical filamentous structures under desiccation-mimicking conditions, and that this molecular assembly was a reversible process. These findings provide insights into the protective mechanisms involved in the anhydrobiosis of tardigrades.

## Results

### Fibril formation of the CAHS1 protein *in vitro*

We first examined molecular structural states of the CAHS1 protein under dehydrating conditions *in vitro*. The dried CAHS1 proteins on a carbon grid were visualized by transmission electron microscopy (TEM), showing that fibril structures were formed (**Figure 1A**). The infrared (IR) spectroscopic analysis showed that the dehydrated CAHS1 protein adopted an α-helical structure (amide I peaks at 1655 cm^-1^ and 1650 cm^-1^), which was reversibly disrupted upon hydration (**Figure 1B and C**). This is consistent with the previous circular dichroism (CD) data, which indicated that the CAHS1 protein adopted helical structures under water-deficient conditions mimicked with trifluoroethanol (19). These data suggest that the CAHS1 proteins underwent fibrilization coupled with the formation of α-helical structures under desiccating conditions.

**Figure 1.**
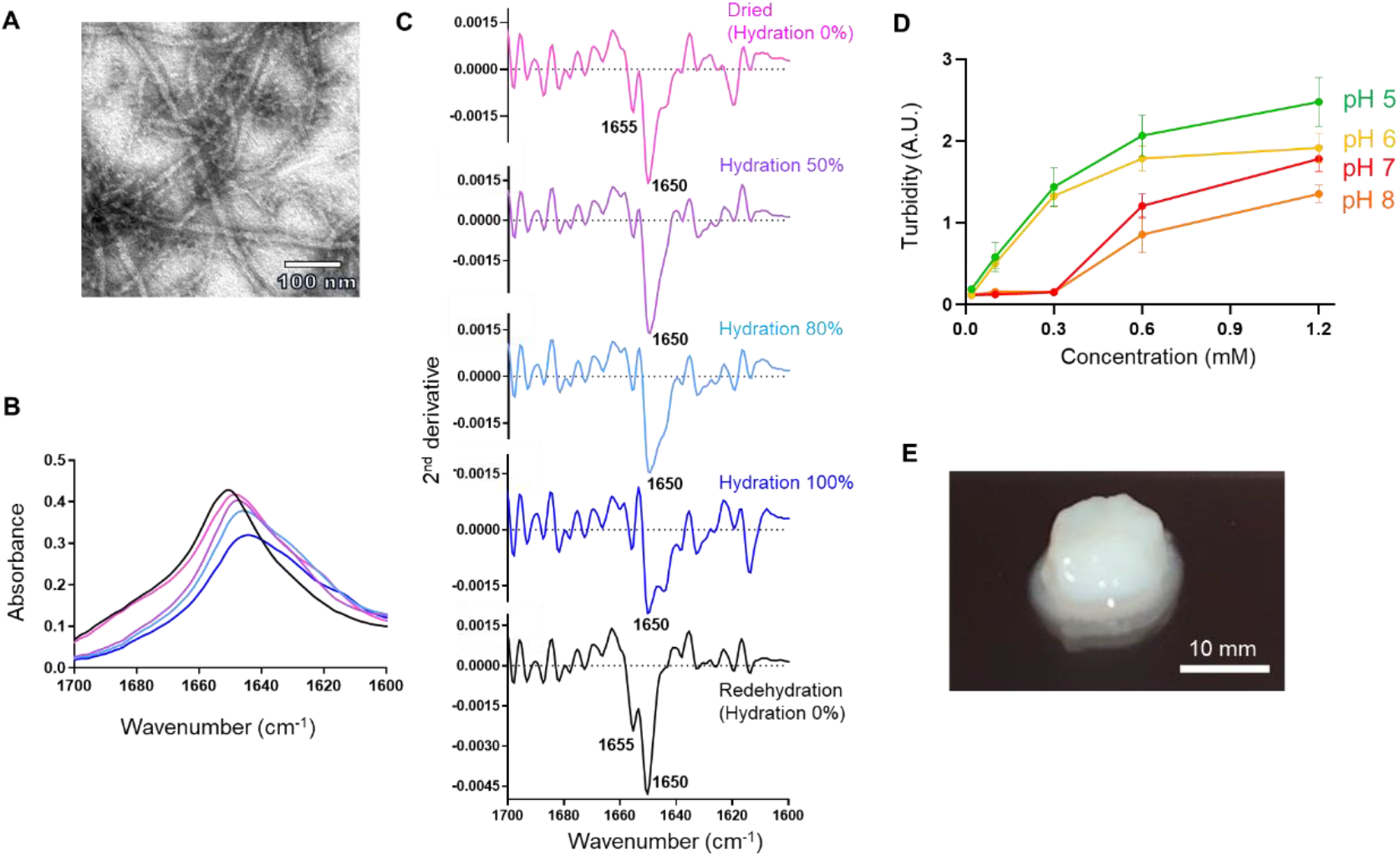
*In vitro* characterization of the fibrous condensation of CAHS1 proteins. (A) TEM image of CAHS1 protein fibrils under dry conditions. (B) Amide I band of the CAHS1 protein and (C) its second derivative under 0 (magenta), 50 (purple), 80 (cyan), and 100% hydration (D_2_O) and dehydration conditions. After 100% hydration (blue), the sample was measured again under redehydration condition (black). (D) Turbidity (absorbance at 595 nm) of the CAHS1 protein at 25°C and pH 5 (green), pH 6 (yellow), pH 7 (red), and pH 8 (orange). The error bars show the standard deviation of three replicates. (E) The CAHS1 protein hydrogel formed in 20 mM potassium phosphate buffer (pH 6.0) at 25°C. The initial protein concentration was 1.2 mM.

For a more detailed understanding of this molecular transition, we characterized the assembly of the CAHS1 protein at different concentrations in aqueous solutions, because the protein concentration is supposed to increase upon drying. The turbidity of CAHS1 protein solutions increased as the CAHS1 protein concentration increased. Thus, the CAHS1 protein was apt to condensate at higher concentrations (**Figure 1D**). In a neutral pH range, the turbidity rose abruptly when the CAH1 protein concentration exceeded 0.3 mM. The CAHS1 protein eventually gelled at a higher concentration (>0.6 mM) (**Figure 1E**) and reversibly turned into a solution upon dilution. These findings indicated that the CAHS1 protein underwent a sol-gel transition in a reversible manner (**Figure 1D and E**).

To determine the structural information of its concentration-dependent conformational changes, we performed nuclear magnetic resonance (NMR) spectroscopic analyses of the CAHS1 protein solution. The ^1^H-^15^N HSQC spectral data obtained under protein-dilute conditions were similar to those previously reported for CAHS proteins from *H. exemplaris* (21), whose structures were interpreted to be intrinsically disordered (**Figure 2A**). We noticed that the HSQC peaks from the CASH1 protein at lower concentrations could be classified into two: extremely narrow and broad ones. We divided the CAHS1 protein from *R. varieornatus* into the N-terminal and C-terminal halves (termed CAHS1-N and CAHS1-C, respectively) and measured their NMR and CD spectra (**Supporting Figure S1A-C**). The results indicate that the CAHS1-N and CAHS1-C fragments adopted an intrinsically disordered and α-helical structures, respectively. Furthermore, the NMR spectrum of the CAHS1 protein exhibited a line-broadening in many peaks from the C-terminal segment, which is a characteristic of molten globule states, suggesting the unstable nature of its C-terminal α-helical structure. As the concentration of the CAHS1 protein increased, these broader peaks disappeared, rendering only the narrower peaks that originated from the N-terminal disordered segments observable (**Figure 2A**). These data indicate that the CAHS1 protein forms condensates through its C-terminal helical region, leaving the remaining regions mobile and disordered.

**Figure 2.**
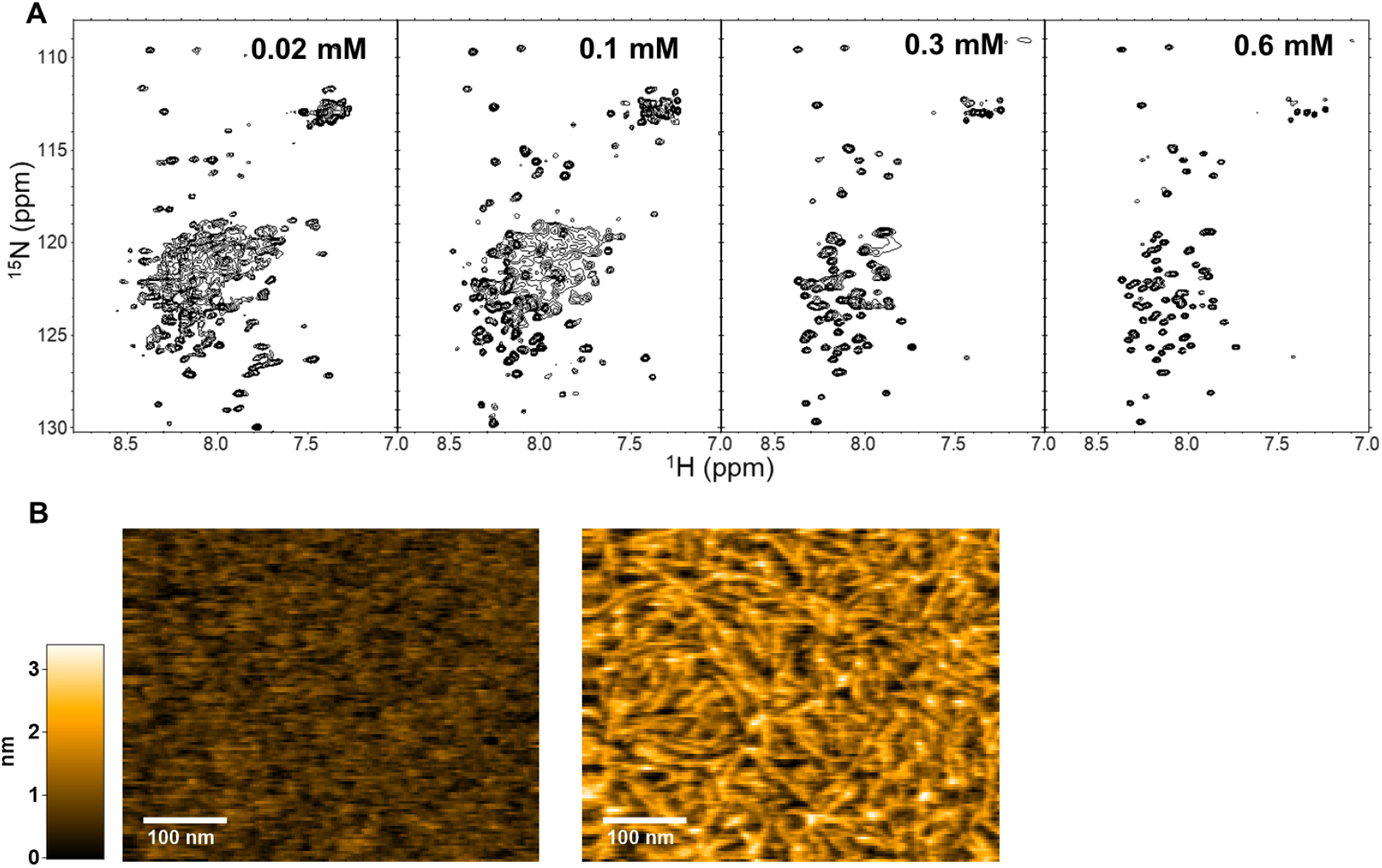
Fibril formation of the CAHS1 protein *in vitro*. (A) ^1^H-^15^N HSQC spectra of the CAHS1 protein at protein concentrations of 0.02, 0.1, 0.3, and 0.6 mM. (B) Typical HS-AFM images of CAHS1 protein fibrils. *Left*: the HS-AFM image of the mica surface with CAHS1 protein at a final concentration of 0.4 μM, about 3 min after injection into the observation solution. *Right*: the HS-AFM image about 2 min after the addition of CAHS1 protein at a final concentration of 3.3 μM.

To observe this sol-gel transition phenomena in a mesoscopic view, we performed a real-time observation of the CAHS1 protein oligomerization using high-speed atomic force microscopy (HS-AFM). Under dilute conditions, the CAHS1 protein was monomeric, consisting of a flexible disordered part and a globular region (**Movie 1**). In contrast, the CAHS1 protein formed fibrils when its concentration was increased (from 0.4 μM to 3.3 μM) (**Figure 2B, Movie 2**). Such fibril formation was observed for CAHS1-C but not CAHS1-N (**Supporting Figure S1D**). Furthermore, the HS-AFM data indicate that the CAHS1 fibrils disappeared upon the addition of 50 mM KCl, indicating the reversible nature of fibril formation (**Supporting Movie S1**). These data indicate that the CAHS1 proteins assembled into fibrous structures through their C-terminal helical regions, leading to gel formation.

### Fibril formation of the CAHS1 protein *in vivo*

The *in vitro* experimental data showed that CAHS1 proteins assembled into fibrils at high concentrations in solution as well as under desiccating conditions. We further examined the potential ability of the CAHS1 protein to form aggregates under molecular crowding condition in cells. In a previous report, the bacterial expression of CAHS proteins from *H. exemplaris* resulted in an improved tolerance of bacterial cells to desiccation (21). Hence, we overexpressed the CAHS1 protein from *R. varieornatus* with an N-terminal FLAG tag (FLAG-CAHS1) in *Escherichia coli* BL21(DE3) as a model system and subjected it to TEM observation. The TEM images visualized the intracellular fibril structures with approximately 10-nm width (**Supporting Figure S2**), confirming that CAHS1 proteins can form fibrils in cells.

We hypothesized that the fibril formation of CAHS1 protein in cells is associated with an improved tolerance against various water stresses, such as dehydration and hyperosmotic treatment. Therefore, we attempted to assess the relationship between the condensation of CAHS1 protein and hyperosmotic stress. For this purpose, the CAHS1 protein with a C-terminal FLAG (CAHS1-FLAG) was overexpressed in HeLa cells, which are stress-sensitive human cells. Immunostaining with anti-FLAG antibody visualized the cytoplasmic distribution of the recombinant CAHS1 protein (**Supporting Figure S3**), which is consistent with a previous study (19). After the hyperosmotic shock with 0.5 M sorbitol, the CAHS1-FLAG proteins were deposited as approximately 650-nm particles. Furthermore, for live imaging, the CAHS1 protein fused to a C-terminal mEGFP protein (CAHS1-mEGFP) was overexpressed in HeLa cells. The fluorescence-labeled CAHS1 proteins were distributed evenly throughout the cytoplasm in the absence of osmotic stresses, but these were rapidly assembled into bright particles by a sorbitol-induced osmotic shock (**Figure 3, Movie 3**). Upon the removal of sorbitol by buffer replacement, the particles disappeared, indicating that the condensation of CAHS1 protein was reversible.

**Figure 3.**
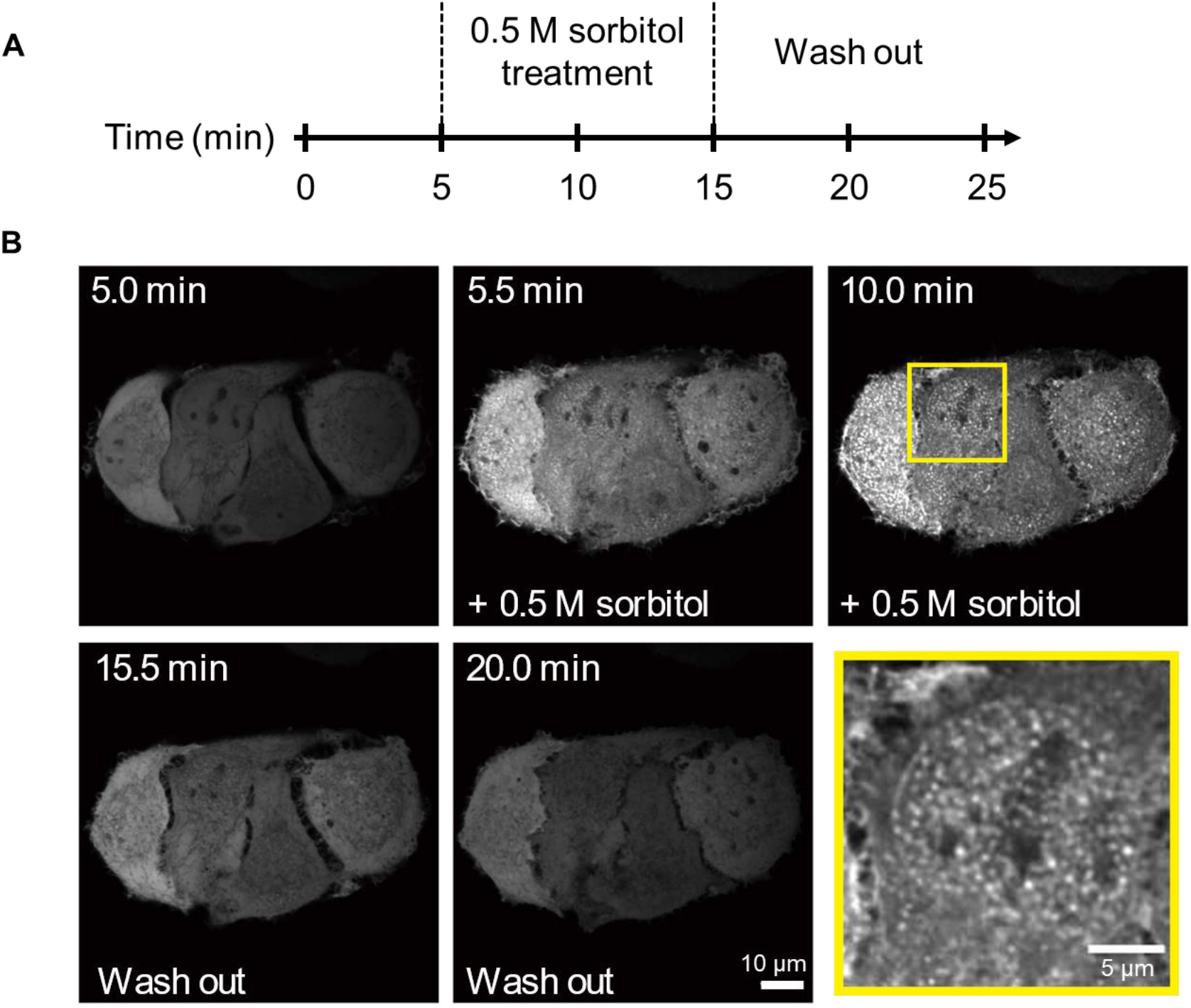
Real-time monitoring of the reversible formation of CAHS1 protein particles. (A) Timeline of time-laps imaging with hyperosmotic shock. (B) Time-laps images of HeLa cells overexpressing the CAHS1-mEGFP proteins. A sorbitol solution was added at 5 min. After 10 min, the cells were washed with a flesh medium using a flow system. An enlarged image of the boxed area enclosed by a yellow line is shown in the lower right.

## Discussion

In this study, the self-assembling property of CAHS1 protein was characterized *in vitro* and *in vivo*. The *in vitro* data indicated that the CAHS1 proteins homo-oligomerized *via* their C-terminal α-helical regions as their concentration increases. The HS-AFM data clearly visualized the fibril formation process of this protein. Furthermore, the *in vivo* observation demonstrated that the overexpressed CAHS1-mEGFP proteins formed condensates in an osmotic shock-dependent manner. At higher protein concentrations, the CAHS1 proteins formed a hydrogel presumably associated with its condensation. These data strongly suggest that, upon drying, the CAHS1 proteins in tardigrade cytosols forms oligomers and eventually undergoes sol-gel transition. The present data also showed that the α-helical fibrous network of the CAHS1 protein can be formed in *E. coli* cells and are maintained even in desiccated conditions. These molecular assembling processes are reversible, depending on solution pH, temperature, and salt concentration.

The multistep phase transition of the CAHS1 protein suggests its multifunctional property. In particular, the CAHS1 fibril formation implies that the anhydrobiotic functions of this protein are distinct from those of trehalose, i.e., water replacement and/or vitrification. Indeed, although the CAHS protein vitrification was hypothesized to be a key mechanism (21), the interpretation of the data leading to such hypothesis has been negated (25). The osmotic shock-induced formation of cytosolic coacervate-like droplets is reminiscent of the stress granules, which are supposed to protect the mRNAs under stress conditions (26). It has recently been reported that a LEA protein from *A. franciscana* forms droplets incorporating positively charged fluorescent protein (27). The present study along with this report suggests that the liquid-liquid phase separation involving the protein assembly promotes the desiccation tolerance through reversible formation of protective compartments for desiccation-sensitive biomolecules. In these compartments, the CAHS1 proteins presumably form a fibrous network with a water-holding capacity that is resistant to desiccation. Moreover, under extremely anhydrobiotic conditions, the CAHS1 network may act as a “dry chaperone,” which not only protects the desiccation-sensitive proteins against denaturation, but also maintains the integrity of other biomolecular complexes, including RNAs and membranes.

We thus propose that, like the LEA proteins, the CAHS1 proteins form cytosolic fibrous condensates, which presumably have variable mechanisms for the desiccation tolerance of tardigrades. There exist a series of LEA-like proteins, including the CAHS-family proteins, which exhibit different cellular distributions and subcellular localizations (19, 20, 22). These proteins may have different assembling properties from those of the CAHS1 proteins, playing distinct roles in a coordinated manner in anhydrobiotic processes. A comprehensive characterization of these proteins is currently underway in our groups.

## Material and methods

### Expression and purification of CAHS1 protein

The plasmid vector encoding the *R. varieornatus* CAHS1 protein was constructed and cloned as a fusion protein with a hexahistidine (His6) tag at the N-terminus using a pET28a vector (Novagen, Merck). CAHS1-N (Met1-Ser96) and CAHS1-C (Pro97-His237) were constructed using standard genetic engineering techniques with a His6 tag at the N-terminus using the pET28a vector (Novagen, Merck). The recombinant CAHS1 protein, CAHS1-N, and CAHS1-C were expressed in the *E. coli* BL21(DE3) strain in Luria Bertani media. For the production of ^15^N-labeled proteins, cells were grown in M9 minimal media containing [^15^N]NH_4_Cl (1 g/L). After sonication, the supernatant was incubated at 90°C for 30 min; then, heat-soluble and -insoluble fractions were separated by centrifugation. The His6-tag fusion proteins were purified with a cOmplete™ His-tag purification resin (Roche, Merck). The fusion proteins were cleaved by incubation with thrombin protease (Sigma-Aldrich) and purified with a Superdex 200 pg (Cytiva) with 10 mM potassium phosphate buffer (pH 7.0).

### NMR measurements

The NMR spectral measurements were made on a Bruker DMX-500 spectrometer equipped with a cryogenic probe. The probe temperature was set to 5°C. ^15^N-labeled CAHS1 protein, CAHS1-N, and CAHS1-C were dissolved at a protein concentration of 0.1 mM in 20 mM potassium phosphate buffer (pH 7.0) containing 5% (v/v) 2H_2_O.

### CD measurements

The CD spectra were measured at 25°C on a JASCO J-720WI apparatus using a 1.0-mm path length quartz cell. The CAHS1 protein, CAHS1-N, and CAHS1-C were dissolved at a protein concentration of 10 μM in 20 mM potassium phosphate buffer (pH 7.0).

### Turbidity measurements

The CAHS1 protein in various conditions were added to a 96-well microplate (Corning, #4442). The optical density of the CAHS1 protein was measured at 25°C at 595 nm by a multi-mode microplate reader (Infinite 200 PRO, TECAN). The CAHS1 protein was dissolved at protein concentrations of 0.02, 0.1, 0.3, 0.6, and 1.1 mM in 20 mM potassium phosphate buffer (pH 8.0, 7.0, 6.0, or 5.0) with or without 150 mM NaCl.

### IR measurements

Attenuated total reflection-Fourier transform infrared measurements were performed using an FTIR spectrometer (VERTEX70, Bruker Optics) equipped with a mercury-cadmium-telluride detector. A 5 μL aliquot of 40 μM CAHS1 proteins (1.1 mg/mL) in ultrapure water was placed on the surface of a diamond ATR crystal with three internal reflection (DurasamplIR II, Smiths Detection) and dried under a gentle stream of N2 gas. After the spectral measurements of dried state of CAHS1 proteins, two 2 μL aliquots of D_2_O or glycerol-OD_3_/D_2_O mixture solution were placed beside the ATR crystal surface and sealed together with the CAHS1 sample by a CaF_2_ window (25-mm diameter and 2-mm thickness) and silicone spacer to control the hydration level of the CAHS1 samples through saturated heavy water vapor. Varying amounts of glycerol-OD_3_ liquid (Sigma, purity 99.0%) were mixed with D_2_O to obtain glycerol/water solutions with 0–50 volume % of glycerol, which enabled the regulation of the saturated vapor pressure in a closed compartment (28). Using D_2_O instead of H_2_O, the amide I band of proteins was observed free from interference with the OH bending mode of water. After 5 min, the spectra were measured at 25°C. Each spectrum was collected with a spectral resolution of 2 cm^-1^ in the 4000–700 cm^-1^ region by averaging 32 scans of interferograms. A background spectrum was recorded with an empty ATR crystal before the measurement of samples. The second derivatives of the spectral data were analyzed by an IGOR Pro software.

### HS-AFM

The HS-AFM images of the CAHS1 proteins were acquired in a tapping mode using a laboratory-built HS-AFM apparatus and a short cantilever (Olympus: BL-AC10;9-μm long, 2-μm wide, and 130-nm thick, 500 KHz-600 kHz resonance frequency in observation buffer, 0.1 N/m spring constant) at 25°C. To observe its monomeric structure, the CAHS1 protein was dissolved at a concentration of 10 nM in 10 mM potassium phosphate buffer (pH 7.4) and deposited on a freshly cleaved mica surface. To observe the fibril formation of the CAHS1 protein at its higher concentrations, the mica substrates with 0.4-μM CAHS1 protein in pure water was imaged, and after about 3 min, the CAHS1 protein solution was added to the solution so that the final concentration was 3.3 μM. The HS-AFM observation of CAHS1-N was performed in the same way as for the full-length CAHS1 protein. For HS-AFM observation of CAHS1-C, the mica substrates with 2-μM CAHS1-C were imaged in 10 mM sodium phosphate buffer (pH 7.4). To investigate the reversibility of fibril formation, 50 mM KCl was added to the CAHS1 fibrils.

### TEM

FLAG-CAHS1 was constructed using standard genetic engineering techniques with an N-terminal FLAG using a pT7-FLAG-1 vector (Sigma-Aldrich, Merck). The bacterial cells of *E. coli* BL21(DE3) strain transformed with FLAG-CAHS1 plasmids were cultured in M9 minimal media at 37°C under constant shaking. Protein expression was induced by adding 0.5 mM isopropyl-β-D-thiogalactopyranoside when the absorbance reached 0.8 at 600 nm. After 4 h, the cells were harvested and washed twice with 50 mM Tris-HCl (pH 8.0) and 150 mM NaCl. The control was run without the FLAG-CAHS1 plasmid.

Cell pellets collected were fixed with 2.5% glutaraldehyde in 10 mM Tris-HCl (pH 8.0) for 30 min at 23°C and collected by centrifugation at 5,000 g for 5 min. The cells were washed with distilled water three times to remove glutaraldehyde and fixed with 2% osmium tetroxide at 23°C for 1 h. After washing with distilled water, fixed cells were stained with 1% uranyl acetate for 20 min. Then, after washing with distilled water, cells were collected by centrifugation at 5,000 g for 3 min. Next, the pellet was embedded in 2% low melting point agar medium (Sigma Aldrich, St. Louis, Missouri, USA), then the cell-containing agar medium was cut into small pieces of approximately 1-2 mm and dehydrated with a stepwise ethanol series (50%, 60%, 70%, 80%, 90%, 95%, and 100%). The sample was infiltrated with QY-1 (Nissin EM Co. Ltd., Tokyo, Japan), embedded in epoxy resin (Durcupan: Sigma-Aldrich, St Louis, Missouri, USA), and polymerized at 60°C for 2 days. Ultrathin sections (approximately 70 nm thick) were prepared with a diamond knife and collected on a Formvar-coated Cu single slot grid. Sections were stained with 2% uranyl acetate and 0.5% lead citrate and observed with a transmission electron microscope (JEM1010, JEOL, Tokyo, Japan) at an accelerated voltage of 80 kV.

### Immunofluorescence measurements

The plasmid DNA expressing the CAHS1-FLAG (pCAGGS-CAHS1-FLAG) protein was transfected into HeLa Italy cells with 293fectin transfection reagent (Thermo Fisher Scientific). The cells were seeded on 4-well glass-base dishes (Greiner) and cultured in Dulbecco’s modified Eagle’s medium (DMEM) supplemented with 10% fetal bovine serum (FBS). After one day, the cells were treated with or without 0.5 M sorbitol in DMEM supplemented with 10% FBS for 5 min. The cells were fixed with 4% paraformaldehyde in phosphate-buffered saline (PBS) for 15 min and were permeabilized with 0.5% Triton X-100 in PBS for 15 min at 25°C. After blocking with 5% goat serum (Abcam, ab7481) in PBS for 1 h at 25°C, the cells were incubated with monoclonal mouse ANTI-FLAG® M2 antibody (Sigma-Aldrich, F1804-50UG, dilution 1:100) overnight at 4°C and then with Alexa Fluor 488-conjugated goat anti-mouse IgG antibody (Thermo Fisher Scientific, A32723, dilution 1:1000) for 1 h in PBS containing 5% goat serum 25°C.

### Time-laps imaging with hyperosmotic shock

HeLa cells were seeded on collagen-coated 35-mm glass-base dishes (IWAKI). The plasmid expressing the CAHS1-mEGFP protein (pCAGGS-CAHS1-mEGFP) was transfected into the HeLa cells with 293fectin transfection reagent (Invitrogen). After 48 h, the medium was replaced with an imaging medium (FluoroBrite DMEM, Thermo Fisher Scientific) supplemented with 1% GlutaMAX (Thermo Fisher Scientific) and 0.1% bovine serum albumin.

For hyperosmotic shock experiments, the cells were initially observed in an imaging medium. Then, 0.5 M sorbitol was added 5 min after starting the observations. After 10 min of incubation in 0.5 M sorbitol, the medium was replaced with an imaging medium, and the cells were observed for another 10 min. The images were acquired every 30 s.

### Fluorescence imaging

The cells were imaged with an IX83 inverted microscope (Olympus, Tokyo) equipped with an sCMOS camera (Prime, Photometrics, Tucson, AZ), a spinning disk confocal unit (CSU-W1; Yokogawa Electric Corporation, Tokyo), and diode lasers at a wavelength of 488 nm. An oil immersion objective lens (UPLXAPO60XO, N.A. 1.42; Olympus) was used. The excitation laser and fluorescence filter settings were as follows: excitation laser, 488 nm; dichroic mirror (DM), 405/488/561 nm; and emission filters, 500–550 nm. During the observation, live cells were incubated with a stage incubator containing 5% CO2 (STXG-IX3WX; Tokai Hit) at 37°C.

## Acknowledgments

We are grateful for Takahiro Bino (NIBB) for help in immunofluorescence measurements and fluorescence imaging. We would like to thank Sachiko Yamada (NIPS) for help in preparation of specimens for TEM experiments. We also thank Yuikiko Isono (IMS) for help in preparation of recombinant proteins. *Chlorella vulgaris* used to feed the tardigrades were partly provided by courtesy of Chlorella Industry Inc.

This work was supported in part by JSPS KAKENHI (Grant Number JP19K07041 to M.Y-U, and JP17H03620 to K.Arakawa), JST CREST (Grant Number JPMJCR17N5 to Y.F.), funds from the Nanotechnology Platform Program (Molecule and Material Synthesis) of the Ministry of Education, Culture, Sports, Science, and Technology (MEXT), Japan, funds from Yamagata Prefectural Government and Tsuruoka City, funds from IMS-IIPA internship program to E.W.G., and funds from ENSCP internship program to V.S.. This work was also supported by the Joint Research of the Exploratory Research Center on Life and Living Systems (ExCELLS) (ExCELLS program No. 18-207, 19-208, and 19-501 to K.Arakawa, No. 18-101 to T.U.), by Instrument Center of IMS, and by the Functional Genomics Facility of the NIBB Core Research Facilities.

## Author contributions

Design of research: M. Y.-U., K. Aoki, K. Arakawa, K. K.

Writing: M. Y.-U., K. Aoki, K. Arakawa, K. K.

Protein expression and purification: M. Y.-U., S. N., V. S., E. G., S. Y., T. S.

NMR, CD, DLS, turbidity: M. Y.-U., S.N., V. S., E. G., T. S.

HS-AFM: H. W., C. G., T. U.

IR: M. Y.-U., S. Y., Y. F.

EM: C. S., K. M.

Fluorescence measurements: S. T., K. Aoki

---

**Movie 1.** HS-AFM video of a monomeric CAHS1 protein. Scan size, 100 nm × 80 nm. Imaging rate: 0.15 s/frame.

**Movie 2.** HS-AFM video of CAHS1 protein fibril formation. Scan size, 500 nm × 500 nm. Imaging rate: 0.3 s/frame (× 5 play).

**Movie 3.** Real-time monitoring of the reversible particle formation of CAHS1-mEGFP protein by confocal fluorescence microscopy.

