## Supplemental Figures for "Desiccation-induced fibrous condensation of CAHS protein from an anhydrobiotic tardigrade"

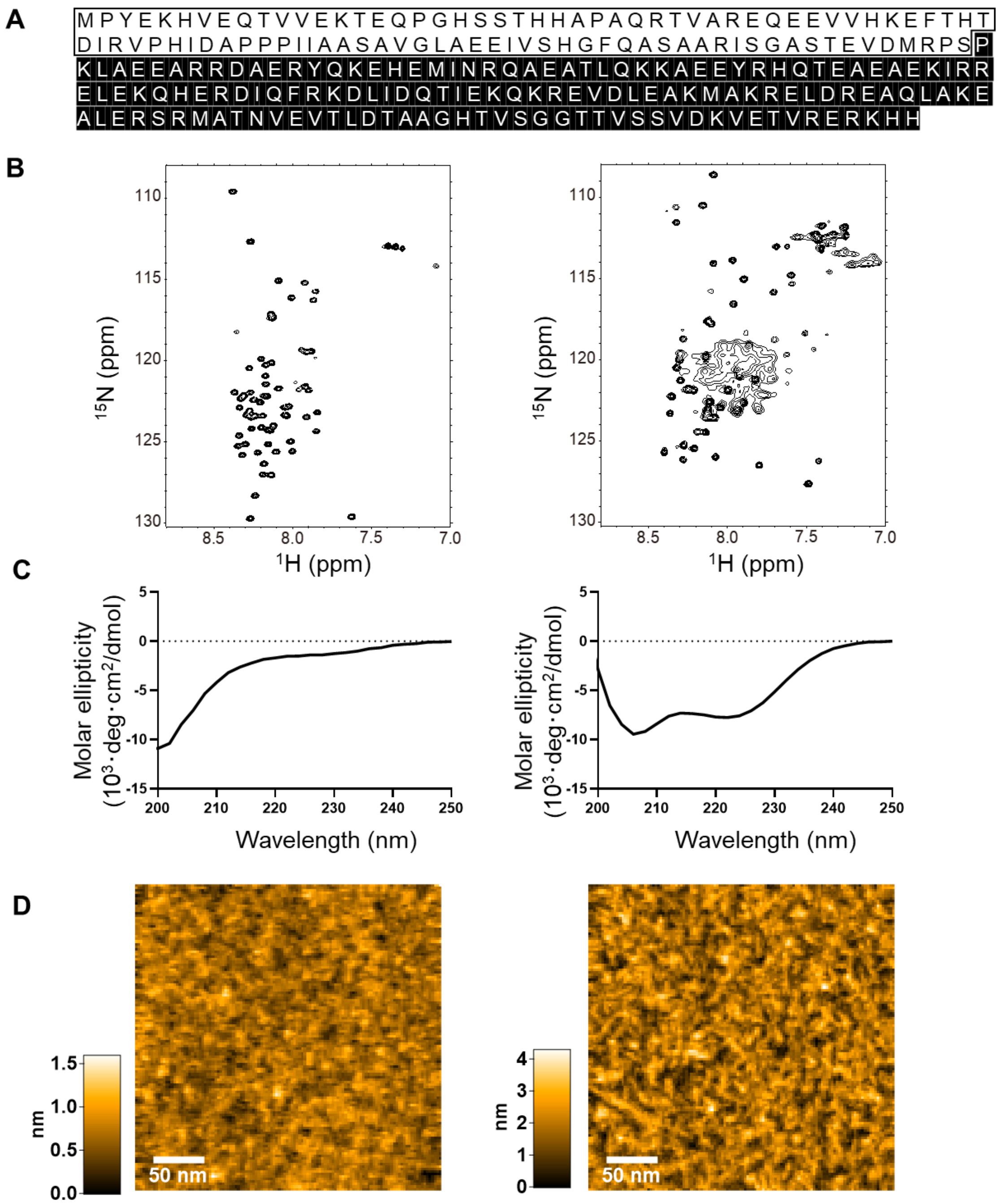


**Figure S1.** ***In vitro* characterization of CAHS1-N and CAHS1-C.** (A) The primary structure of CAHS1 protein with white and black boxes that indicate CAHS1-N and CAHS1-C, respectively. (B) $^{1}$H-$^{15}$N HSQC spectra of CAHS1-N (left) and CAHS1-C (right). (C) CD spectra of CAHS1-N (left) and CAHS1-C (right). (D) Typical HS-AFM images of CAHS1-N (left) and CAHS1-C (right).

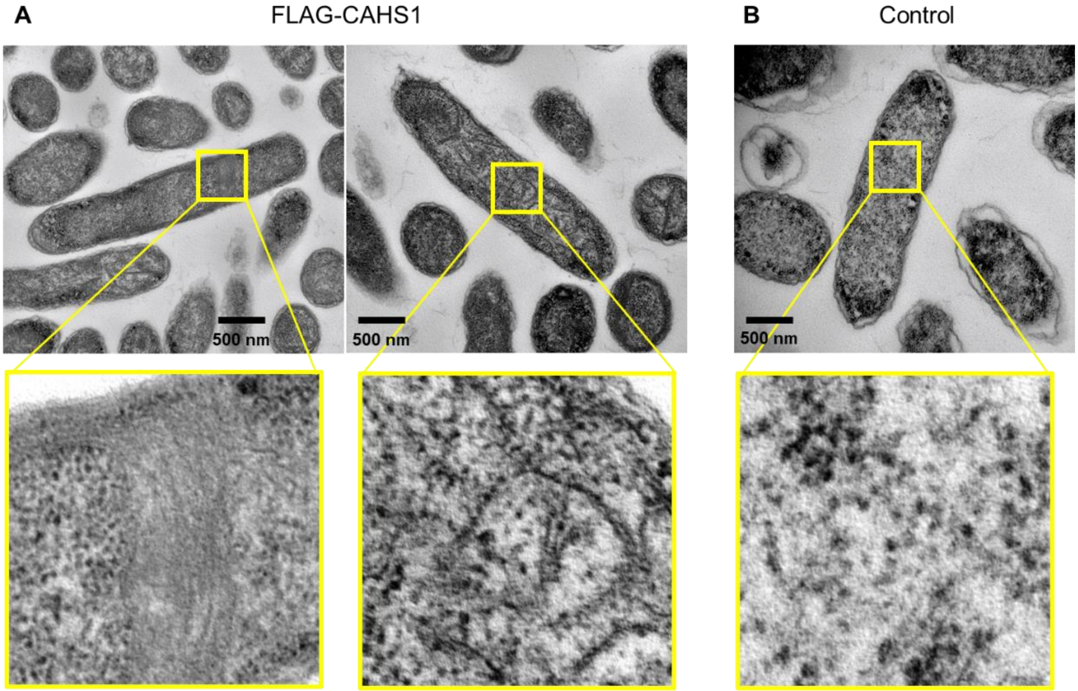


**Figure S2. Fibril formation of the CAHS1 protein in *E. coli*.** Cross-sectional TEM images of *E. coli* (A) with and (B) without overexpressed FALG-CAHS1 proteins (top). Enlarged views of the boxed areas of TEM images (bottom).

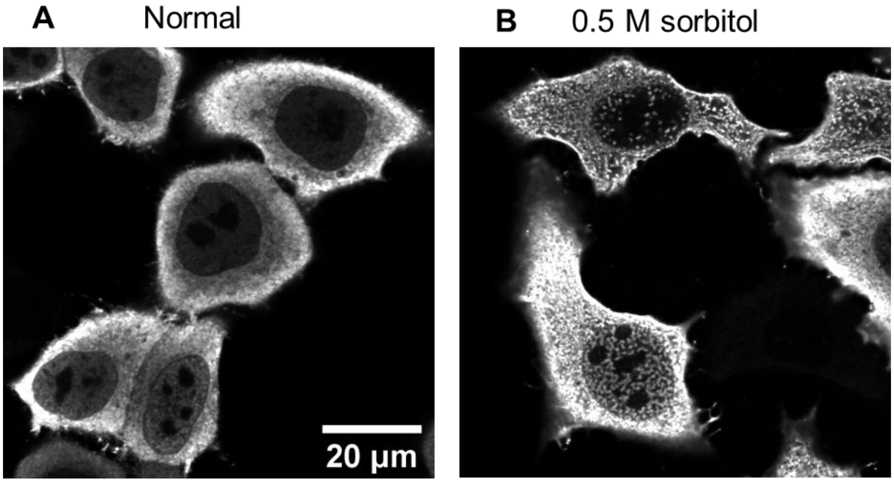


**Figure S3. Osmotic stress induced the CAHS1 protein particles in HeLa cells.** The HeLa cells overexpressing the CAHS1-FLAG proteins were exposed to (A) mock or (B) hyperosmotic medium (0.5 M sorbitol) and stained with anti-FLAG antibody.

**Movie S1.** HS-AFM observation of the disassembly of CAHS1 protein fibrils upon adding 50 mM KCl. Scan size, 200 nm × 160 nm. Imaging rate: 0.2 s/frame (× 2 play).
