## Supplementary figures and images for "Desiccation-induced fibrous condensation of CAHS protein from an anhydrobiotic tardigrade"

### Movie 3.gif

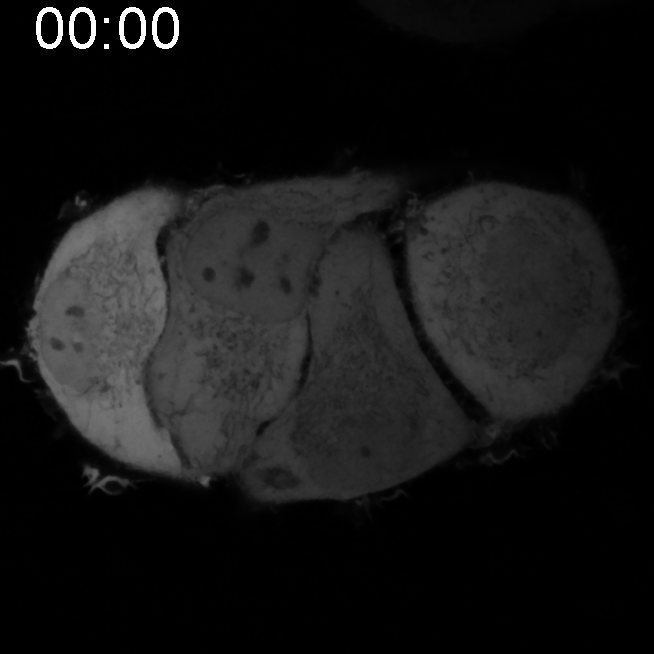
